# DiGeST: Distributed Computing for Scalable Gene and Variant Ranking with Hadoop/Spark

**DOI:** 10.1101/168633

**Authors:** Yann-Aël Le Borgne, Raphaël Helaers, Tom Lenaerts, Marc Abramowicz, Guillaume Smits, Gianluca Bontempi

## Abstract

**Background:** The advent of next-generation sequencing technologies has opened new avenues for clinical genomics research. In particular, as sequencing costs continue to decrease, an ever-growing number of clinical genomics institutes now rely on DNA sequencing studies at varying scales - genome, exome, mendeliome - for uncovering disease-associated variants or genes, in both rare and non-rare diseases.

A common methodology for identifying such variants or genes is to rely on genetic association studies (GAS), that test whether allele or genotype frequencies differ between two groups of individuals, usually diseased subjects and healthy controls. Current bioinformatics tools for performing GAS are designed to run on standalone machines, and do not scale well with the increasing size of study designs and the search for multi-locus genetic associations. More efficient distributed and scalable data analysis solutions are needed to address this challenge.

**Results:** We developed a Big Data solution stack for distributing computations in genetic association studies, that address both single and multi-locus associations. The proposed stack, called DiGeST (Distributed Gene/variant Scoring Tool) is divided in two main components: a Hadoop/Spark high-performance computing back-end for efficient data storage and distributed computing, and a Web front-end providing users with a rich set of options to filter, compare and explore exome data from different sample populations. Using exome data from the 1000 Genomes Project, we show that our distributed implementation smoothly scales with computing resources. We make the resulting software stack Open-Source, and provide virtualisation scripts to run the complete environment both on standalone machine or Hadoop-based cluster.

**Conclusions:** Hadoop/Spark provides a powerful and well-suited distributed computing framework for genetic association studies. Our work illustrates the flexibility, ease of use and scalability of the framework, and more generally advocates for its wider adoption in bioinformatics pipelines.

## Background

The goal of most DNA sequencing studies is to identify causal single-nucleotide variations (SNVs) in patients with Mendelian diseases, and more recently combinations of variants in digenic or oligogenic diseases [6, 17, 15]. There however exists millions of mutations in any individual genome, and identifying which ones are disease-causing remains a largely open problem. A common approach to tackle this issue is to rely on Genetic Association Studies (GAS), which are based on the principle that genotypes can be compared directly, using case-control designs [31, 34, 36, 10].

Case-control designs for GAS generally involve a set of *M* variants *v_i_*, 1 ≤ *i* ≤ *M*, and a set of *N* samples *s_j_*, 1 ≤ *j* ≤ *N*. Samples are divided in two groups 𝓖_0_ and 𝓖_1_, for the control and case groups, respectively. Then, an *M*-by-*N genotype* matrix is expressed as **G** = {*z_i,j_*}_1≤*i*≤*M*,1≤*j*≤*N*_ where *z*_*i*,*j*_ ∈ {0,1, 2} is the zygosity of sample *s_j_* for variant *v_i_*, and the values {0,1, 2} code for homozygous reference, heterozygous, and homozygous alternative, respectively. An example of genotype matrix is given in Fig. 1 for a set of 7 samples and 5 variants over 2 genes.

**Figure 1:**
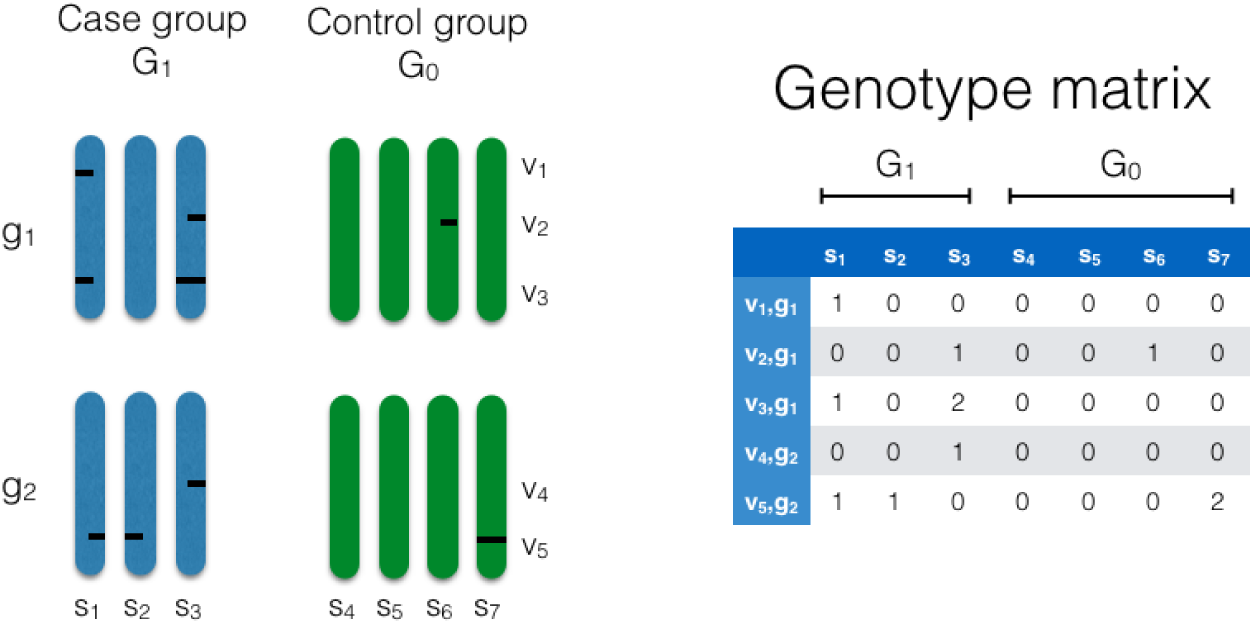
Case/control genotype matrix:

Given a genotype matrix, a GAS aims at finding the sequencing regions that most differ between the case and control groups. This is generally a two step process where, for any given region of interest (for example, variant, gene, or set of genes), the difference between the case and control genotypic data is first quantified by a score and some statistical measure of significance. The scores are then ordered by decreasing order of importance, so that genomic regions of highest difference between the two groups are identified [31, 32, 4, 65].

The standard scoring approach in GAS consists in summing, for each variant, the allele counts for both the case and control groups, and to compute a standard Fisher’s test or χ^2^ statistic for assessing the significance of the difference [31]. Many variations of this baseline approach have been designed. In particular, methods such as CAST [48], CMC [35], WSS [40], KBAC [37], SKAT [70], RareCover (RC) [2] and others [57, 50, 49, 22] have been proposed to ‘collapse’ or aggregate sets of variants within a region, which is usually a gene, in order to increase the statistical power of the tests. Other efforts have been targeted at further extending traditional GAS to gene-gene interaction studies (Genetic Interaction Studies - GIS) [4, 41, 9, 66, 45, 53, 39], in order to address phenomena such as epistasis, now widely accepted as an important contributor to genetic variation in complex diseases.

Score statistics computations often require substantial computational resources. This computational burden becomes especially significant when statistical tests rely on permutations, or when interactions between loci (GIS) are involved. As an example, computing all pairwise scores for a set of one million variants requires on the order of a trillion comparisons. The standard tools for GAS (PLINK [58], VariantTools [67], SnpSift [5], R Bioconductor [16] or AssotesteR [23], gNOME [31]) are however designed to run on standalone machines, and do not scale well as the size and complexity of genetic study designs increase.

Few solutions have been designed to address this challenge, that mostly rely on dedicated hardware devices such as Graphical Processing Units (GPUs) [72, 71, 25, 18], or Field-Programmable Gate Array (FPGAs) [68, 20]. While these solutions greatly speed up computation times, their use is in practice hindered by the need to acquire specialised and expensive hardware, whose programming is based on low-level and difficult to debug programming languages.

In this paper, we argue that the recently developed Spark cluster computing framework, coupled with a Hadoop cluster back-end, provides a well-suited distributed environment for performing large scale genetic association studies. Besides providing a fast and scalable computing framework, Spark features a rich and high-level application programming interface (API) particularly well-suited for manipulating data such as variant data. On the other hand, the Hadoop framework is designed to operate on off-the-shelf computer hardware, provides a distributed and fault tolerant data storage back-end (Hadoop Distributed File System - HDFS), and a robust resource management platform for parallelising Spark jobs.

The contributions of this paper are several. First, we show that Spark can be used to efficiently extract variant data from a distributed database using standard SQL syntax, and to generate genotype matrices as distributed collections of objects. We then detail how scoring can be performed in a distributed way, for genomic regions ranging from variant to genes to pairs or variants or genes. Third, we experimentally show that the proposed workflow, called DiGeST (Distributed Gene/Variant Scoring tool), efficiently scales to case control designs involving thousands of samples and billions of scorings, and effectively distributes computations on the available computing resources. Finally, we provide a Web based front end to interact with DiGeST, allowing a user to easily define filtering criteria, create control and case groups, and explore scoring results. All the tools and data are open source and available from [30].

The paper is structured as follows. We first outline the DiGeST workflow, and describe its main components: data filtering, genotype matrix creation, and scoring. We then present the command line and Web interfaces to run DiGeST. Finally, we illustrate its scalability on datasets involving millions of variants and thousands of samples, and discuss the benefits and limits of Hadoop/Spark for genetic association studies.

## Implementation

DiGeST performs scoring in two stages. First, variants of interest are extracted (filtered) from a variant database, and grouped in an intermediary genotype matrix whose entries are the genotypes of all filtered variant/sample pairs. Second, scoring is applied on the genotype matrix, which returns a sorted scoring matrix containing, for each region of interest (variant, gene, pair of variants/genes), the score together with additional statistics (group scores, p-values, …). Both stages are run in a distributed way thanks to the Spark/Hadoop computing frameworks. The workflow is summarised in Fig. 2, and detailed in the following sections.

**Figure 2:**
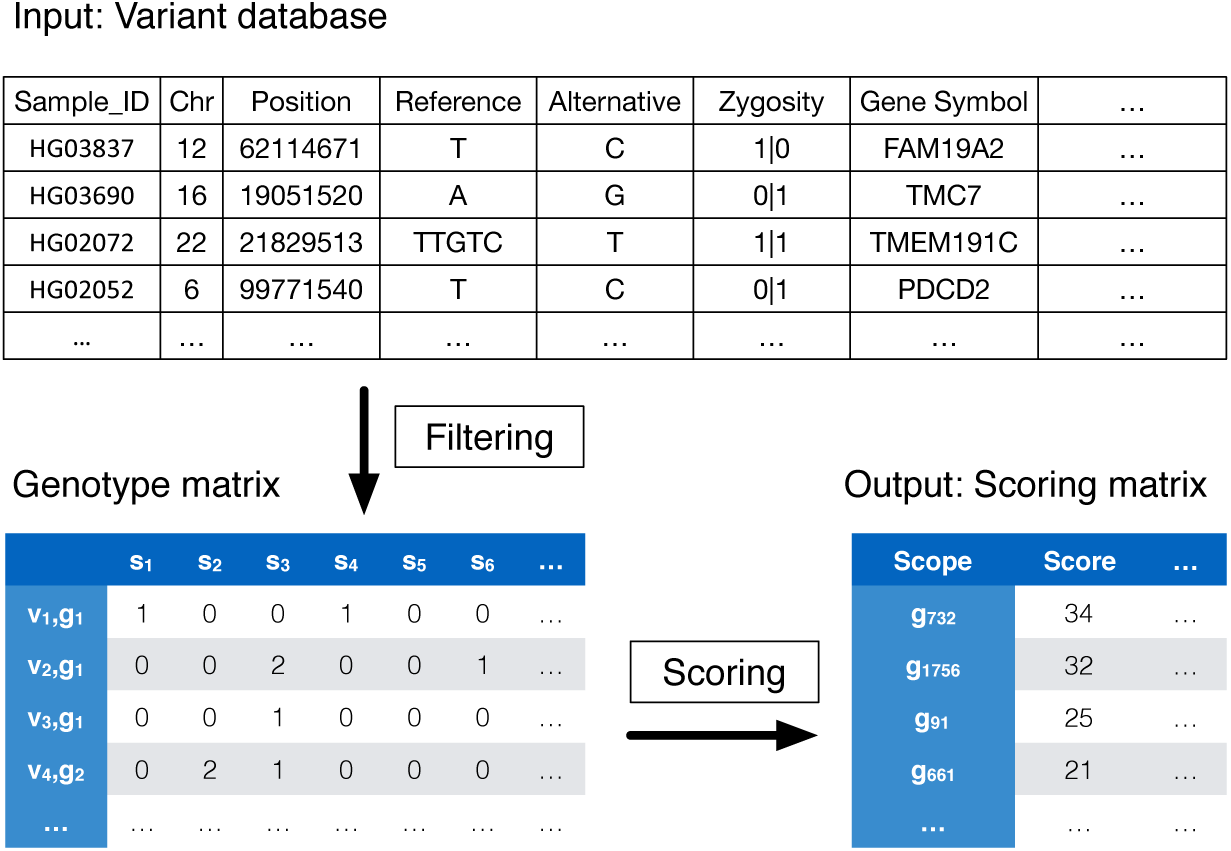
DiGeST Workflow: The variant database, the genotype matrix and the scoring matrix are the key data structures in DiGeST. The two main processing stages (filtering and scoring) are performed using the Spark distributed computing framework.

### Overview of DiGeST workflow and data structures

#### Variant database

The variant database serves as input to the DiGeST workflow, and stores variant data as a single flat table similar to the standard VCF (Variant Call Format) structure [12]. A minimum of seven fields are required for downstream processing with DiGeST. The first field contains the sample identifier. The next four fields (chromosome, position, reference, alternative) uniquely identify the variant by its position and DNA sequence change. The sixth field contains the zygosity of the sample for the given variant, and is represented by 2 (homozygous alternative), 1 (heterozygous alternative) or 0 (homozygous reference). Note that these first six fields are also part of the required fields of VCF files. A seventh column additionally contains the gene symbol, following the HGNC gene nomenclature [19], which is needed by DiGeST for performing ranking along genes.

The database may contain additional fields, which can be used during the filtering stage. Such fields can include, for example, annotations from genomic databases such as 1000 Genomes Project [8], dbSNP [63], dbNSFP [38], SnpEFF [13], that can be obtained using standard annotation software, see [55] for a review.

Given that a single individual genome has 3-4 millions variants, the variant database can become a very large data structure when thousands of samples are included. DiGeST relies on the Apache Parquet format [46, 42] for storing the variant database. Apache Parquet is a columnar data storage format designed to support very efficient compression and encoding schemes. In particular, Parquet allows gzip and snappy compression. While gzip has better compression accuracy (about twice as much as snappy), it is much slower to decompress and compress (about 5 times). As the storage costs continue to decrease, snappy usually provides a better option by significantly reducing the data processing time, allowing end-users to reach their results faster. Besides compression, Parquet also allows files to be split and stored on a distributed file system, and data to be queried from the files using SQL like syntax. Such properties make the format much more suitable for variant filtering than the CSV or VCF formats.

#### Filtering stage

The filtering is the first processing stage of the DiGeST workflow, and consists in extracting variants of interest from the variant database, and creating the genotype matrix as a resilient distributed dataset (RDD). RDDs are Spark’s underlying distributed data structures, split in *partitions*, which allow data to be stored and processed in a distributed manner. They consist in collections of items, usually tuples or (*key*, *value*) pairs. The sequence of transformations from the database extraction to the genotype matrix creation is summarised in Fig. 3.

**Figure 3:**
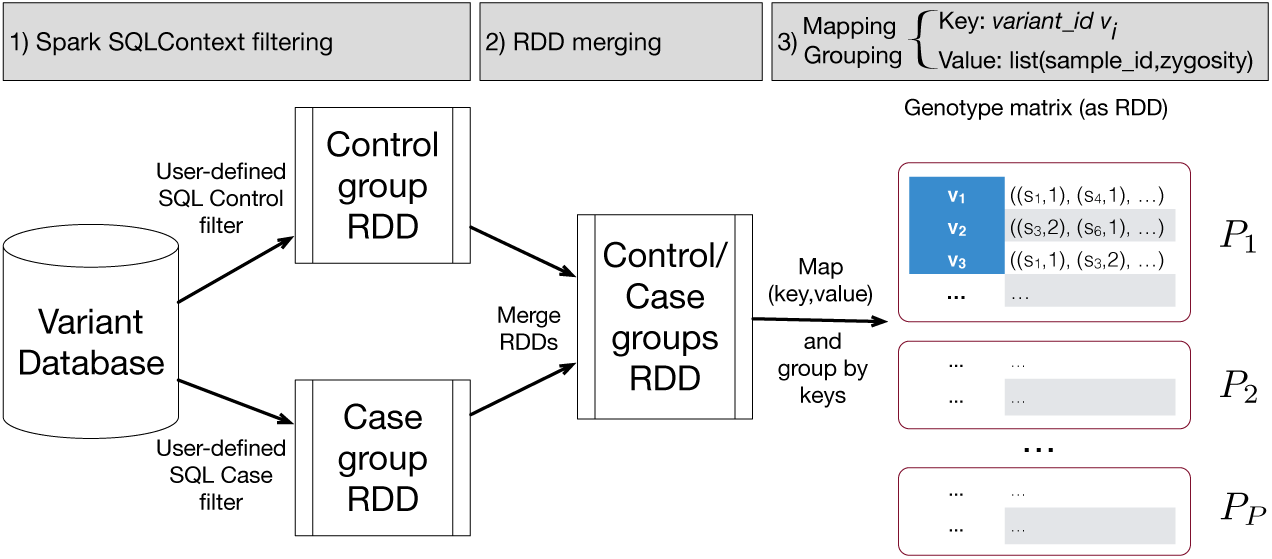
Filtering stage: 1) Spark SQLContext filtering. Variant data are extracted from the variant database using a standard SQL syntax, and stored in RDDs (resilient distributed datasets) for both the control and case groups. 2) RDDs merging. The two RDDs are merged. 3) Mapping. The RDD is rearranged using variants IDs as keys, forming as a result the (sparse) genotype matrix.

The filters are user-defined sets of conditions aimed at selecting variants and samples to include in the control and case groups. They are expressed using a standard SQL syntax, with conditions on the set of fields (columns) available in the variant database. Queries taken by DiGeST to create the case and control groups are typically of the form

~~~
SELECT Sample_ID,Chr,Position,Reference,Alternative,
       Zygosity,Gene_Symbol
FROM variantDatabase
WHERE Sample_ID IN (…)
AND Gene_Symbol IN (…)
AND …
~~~

The set of samples to include in the case and control group is defined by conditioning the *Sample*_*ID* field. Additional conditions can be specified by extending the SQL *WHERE* clause. These may include, for example, conditions on gene symbols to restrict the variant subset to a gene panel, or conditions on quality control or variant deleteriousness annotations if these are available.

SQL queries for the control and case groups are executed using the Spark `SQLContext` object, which provides the entry point to all Spark SQL functionalities. The `SQLContext` returns two RDDs which, after merging, form a unique RDD that contains all variants from the two groups as a distributed collection of tuples (*Sample*_*ID*, *chr*, *pos*, *ref*, *alt*, *zygosity*, *Gene_Symbol*). Finally, in order to group variant data by variant IDs, these tuples are rearranged in (*key*, *value*) pairs, where the *key* consists in a first tuple (*Gene_Symbol*, *chr*, *pos*, *ref*, *alt*) that uniquely identifies a variant, and the *value* is a second tuple (*Sample*_*ID*, *zygosity*) that identifies the zygosity value for a given sample and variant in the genotype matrix. The transformation is done by applying a `createKey_VariantGene` (*variantTuple*) function, see Pseudocode 1.

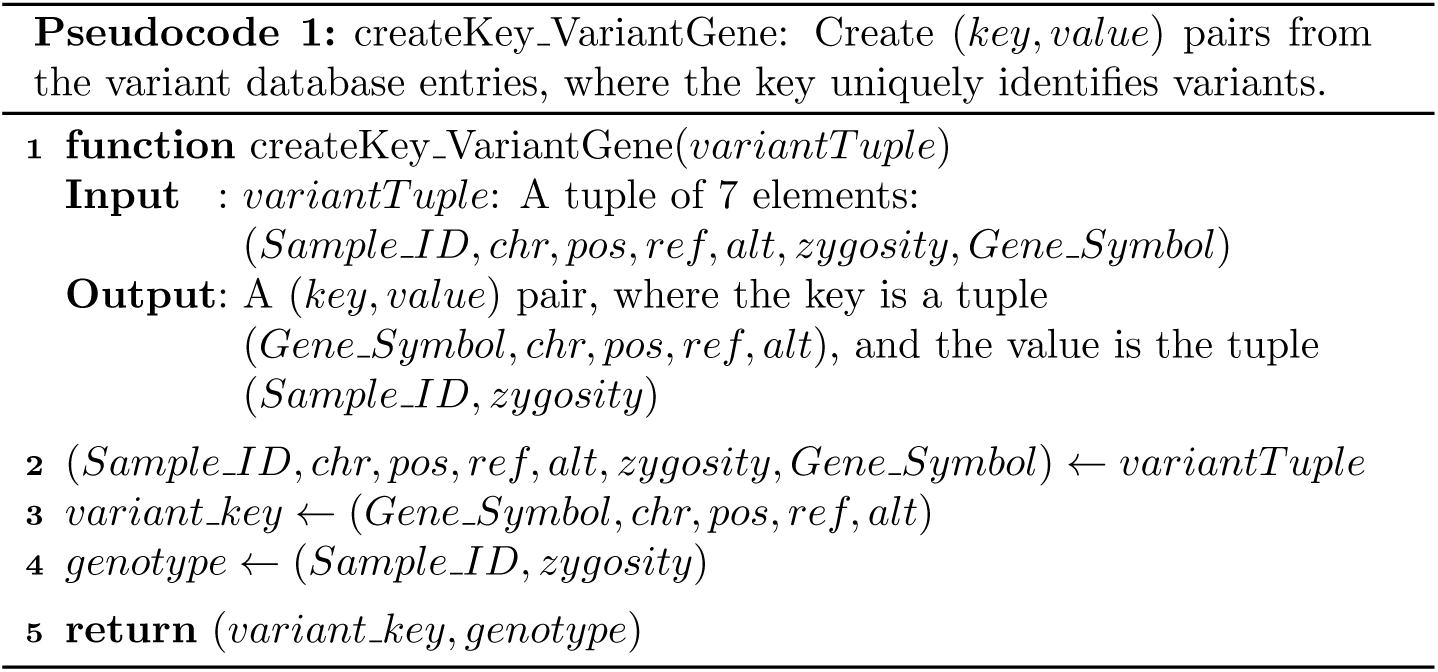

Spark’s application programming interface (API) provides all the high level operators needed to express the transformations summarised in Fig. 3. SQL queries are executed using the `sqlContext` object. Resulting RDDs are then merged using the `union` operator, transformed in (*key*, *value*) pairs using the `map` operator, and grouped by variants using the `groupByKey` operator, as summarised in Pseudocode 2. Since most values in a genotype matrix are sparse, only entries coding for heterozygous and homozygous alternatives (1 or 2, respectively) are effectively stored.

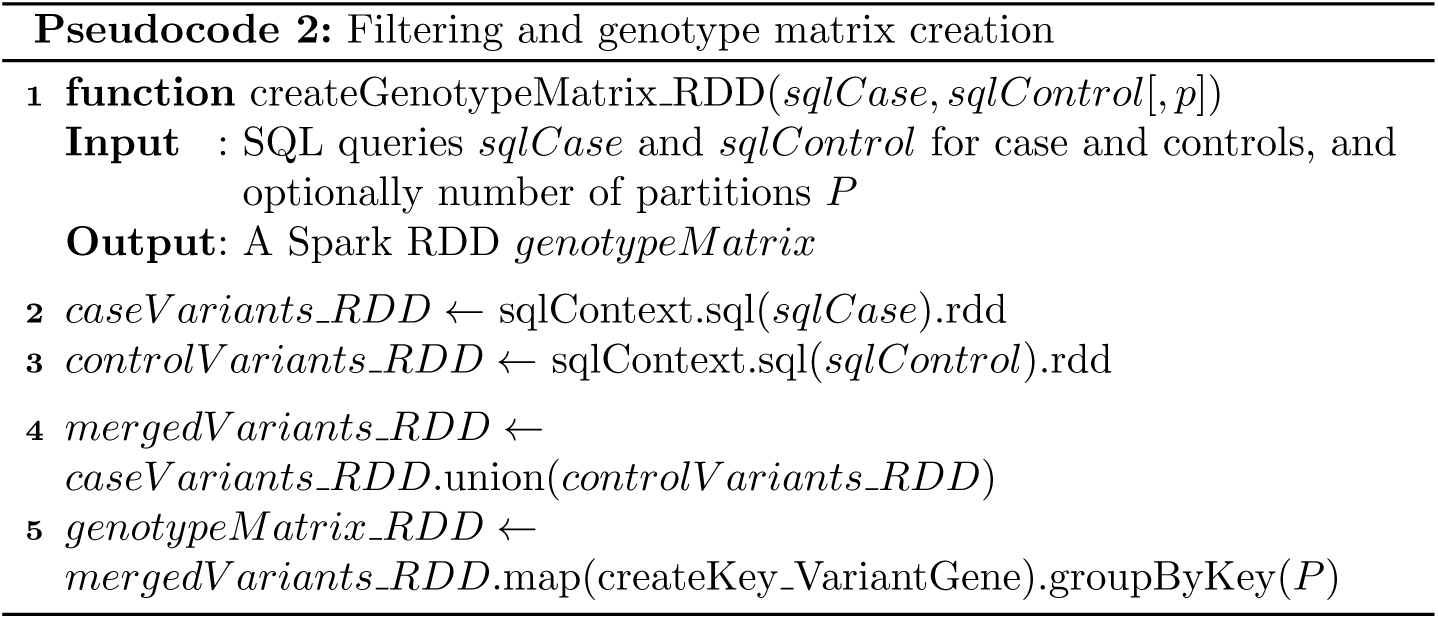

The number of partitions in which the RDD is split when extracting the data from the Parquet database is by default determined by Spark, and depends on the block size of the file system (typically 256MB on HDFS file systems). It may be desirable to increase the number of partitions for better parallelisation, and this can be specified as an optional argument *P* during the grouping operation (Pseudocode 2, line 5).

#### Scoring stage

The scoring stage essentially consists in restructuring the genotype matrix so that the genomic regions to score are grouped together (e.g., genes, pair of variants…), and in applying the scoring function to each group. We outline in the following how Spark can efficiently perform such groupings and compute corresponding scores in a distributed way, for variant, genes and 2–way interactions (pairs of variants/genes). We then show that the proposed approaches can be generalised to the scoring of larger genomic regions (for example sets of genes), and *k*–way interactions.

##### 1) Variant scoring

The Spark implementation for variant scoring is almost straightforward, since the genotype matrix groups genotype data by variant keys. Let us denote by *S*(*P_i_*) a scoring function that takes as input a partition *P_i_* of the genotype matrix, and returns a list of (*variant*_*key*, (*score*, *p*_*value*)) tuples, one for each *variant*_*key* in *P_i_*. An example of Pseudocode for such a function is given in Pseudocode 3, which implements the basic allelic sum score together with a Fisher’s test for statistical significance [31]. Global variables are used to provide the IDs of samples belonging to the case and control groups.

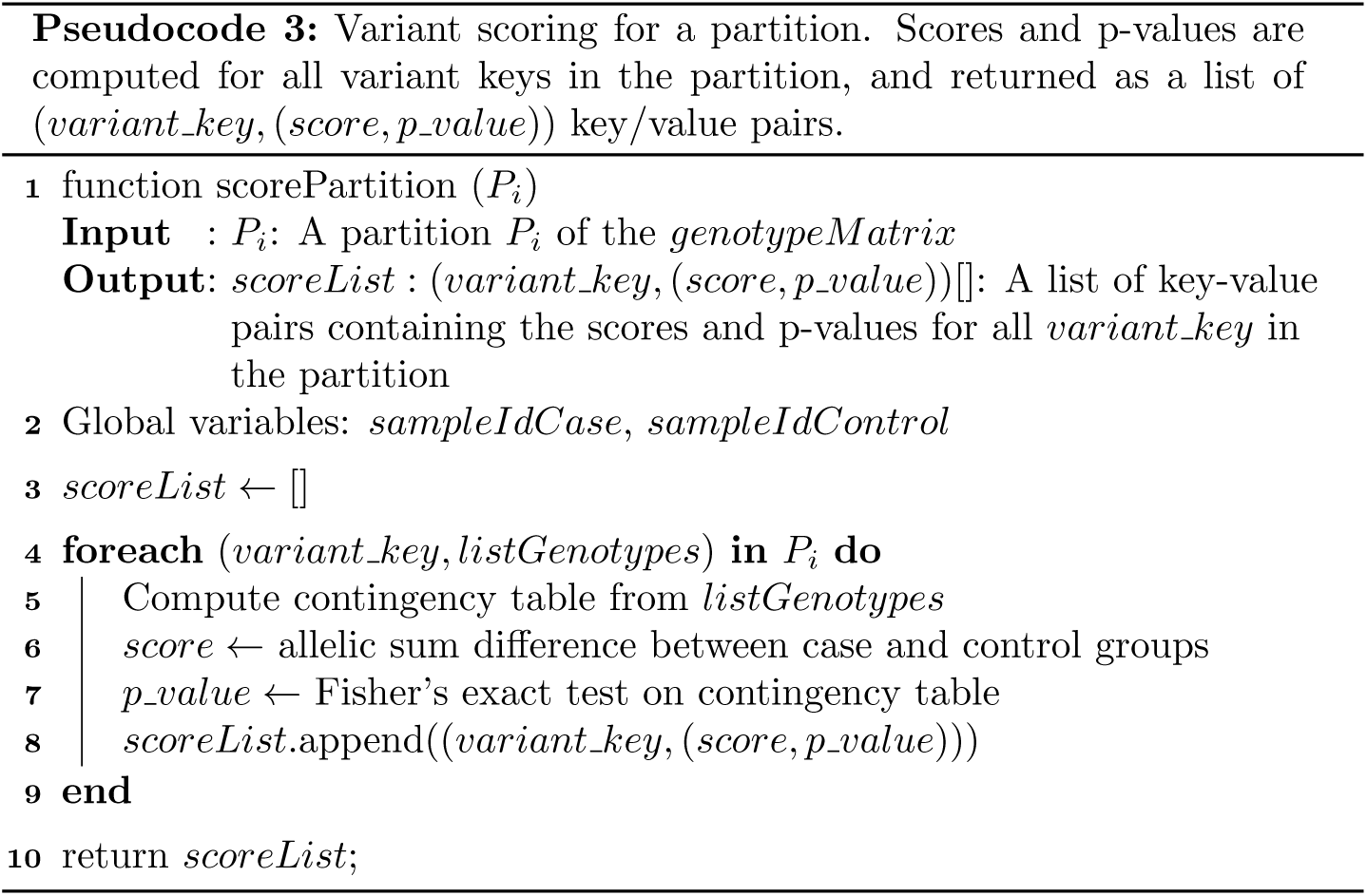

The overall scoring is distributed by applying the scoring function *S*(*P_i_*) to each partition *P_i_* of the distributed genotype matrix by means of the Spark **map** operator. Scores are then sorted the using the `takeOrdered` operator. Finally, sorted variant keys, together with the scores and p-values, are retrieved as a table in the form of a Spark *dataframe.* A high-level summary of the Spark processing pipeline is provided in Fig. 4.

**Figure 4:**
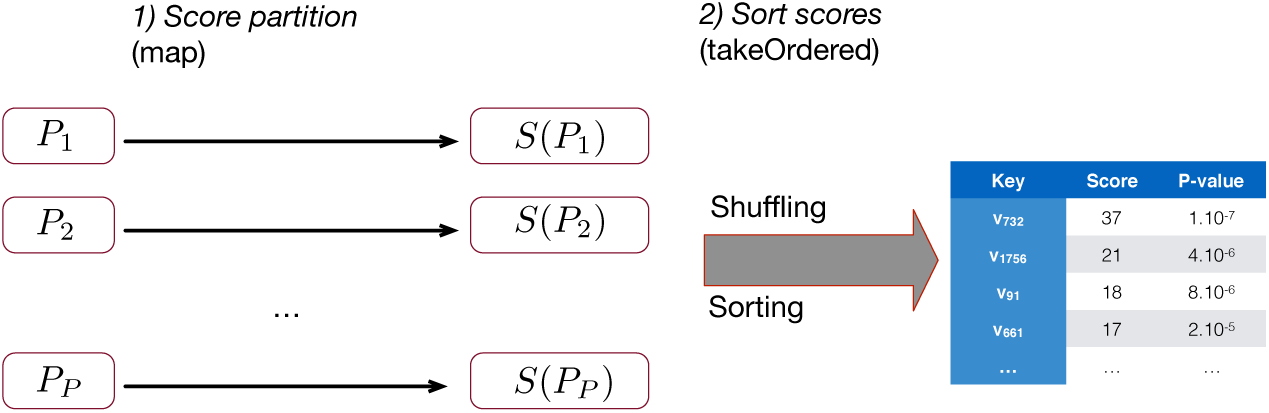
Variant scoring. Within each partition, tuples (variant_key,list(SampleJd, zygosity)) are scored and transformed in tuples (variant_key,(score, p_value)). Variant keys whose scores are the highest are then returned as a Spark *dataframe* using the distributed `takeOrdered` operator.

It is worth noting that the first stage (scoring) does not require any exchange of data (shuffling), while the sorting does, which is represented in the diagram using a thick arrow.

##### 2) Gene scoring

The workflow for gene scoring is essentially the same as for variant scoring, with an added preprocessing step that changes the grouping of the genotype matrix in order to group variant keys belonging to the same gene. The change of grouping is done by using the Spark **groupBy** operator, which takes care of reorganising the genotype matrix along gene keys. The resulting workflow is illustrated in Fig. 5. The change of grouping may require data to be shuffled, which is represented by the thick arrow between the grouping and scoring steps.

**Figure 5:**
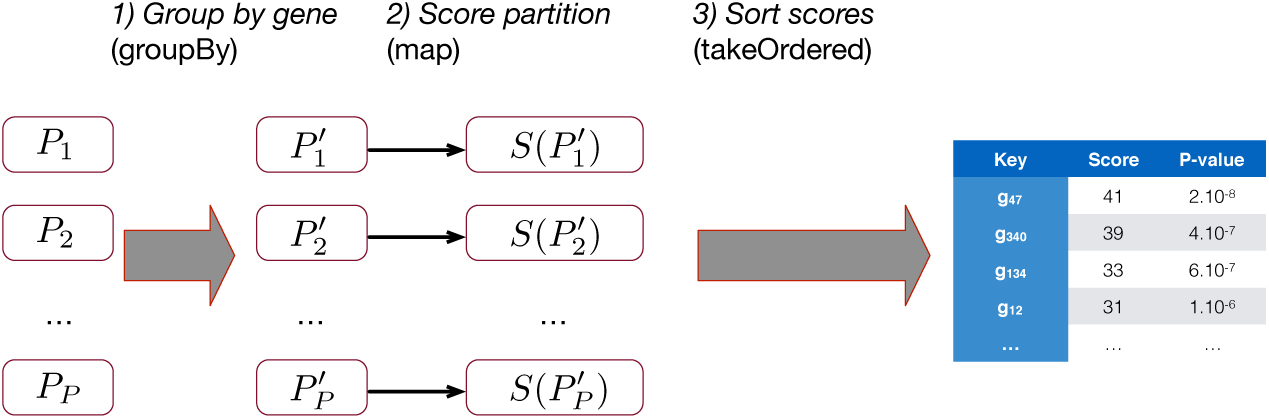
Gene scoring: An additional preprocessing step groups the genotype data using gene as keys.

The scoring of the partitions follows the same logic as for the scoring of variants. The Pseudocode 3 only needs to be adapted in order to perform scoring along gene keys instead of variant keys.

##### 3) Pair of variants scoring

The scoring of pairs of variants requires the genotype matrix partitions to be paired in order to compute the scores for any pair of variants. The set of *P* partitions is therefore first transformed in a set of 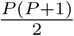 paired partitions, each of which contains a pair of partitions (*P_i_*,*P_j_*),*j* ≥ *i*. An efficient approach to perform this pairing is to build a list *listPairsToCreate* of index pairs (*i*, *j*), for all the pairs of partitions that need to be combined. Index pairs are then used as keys, and partitions *P_i_* are mapped to all index pairs that contain index *i*, see Pseudocode 4.

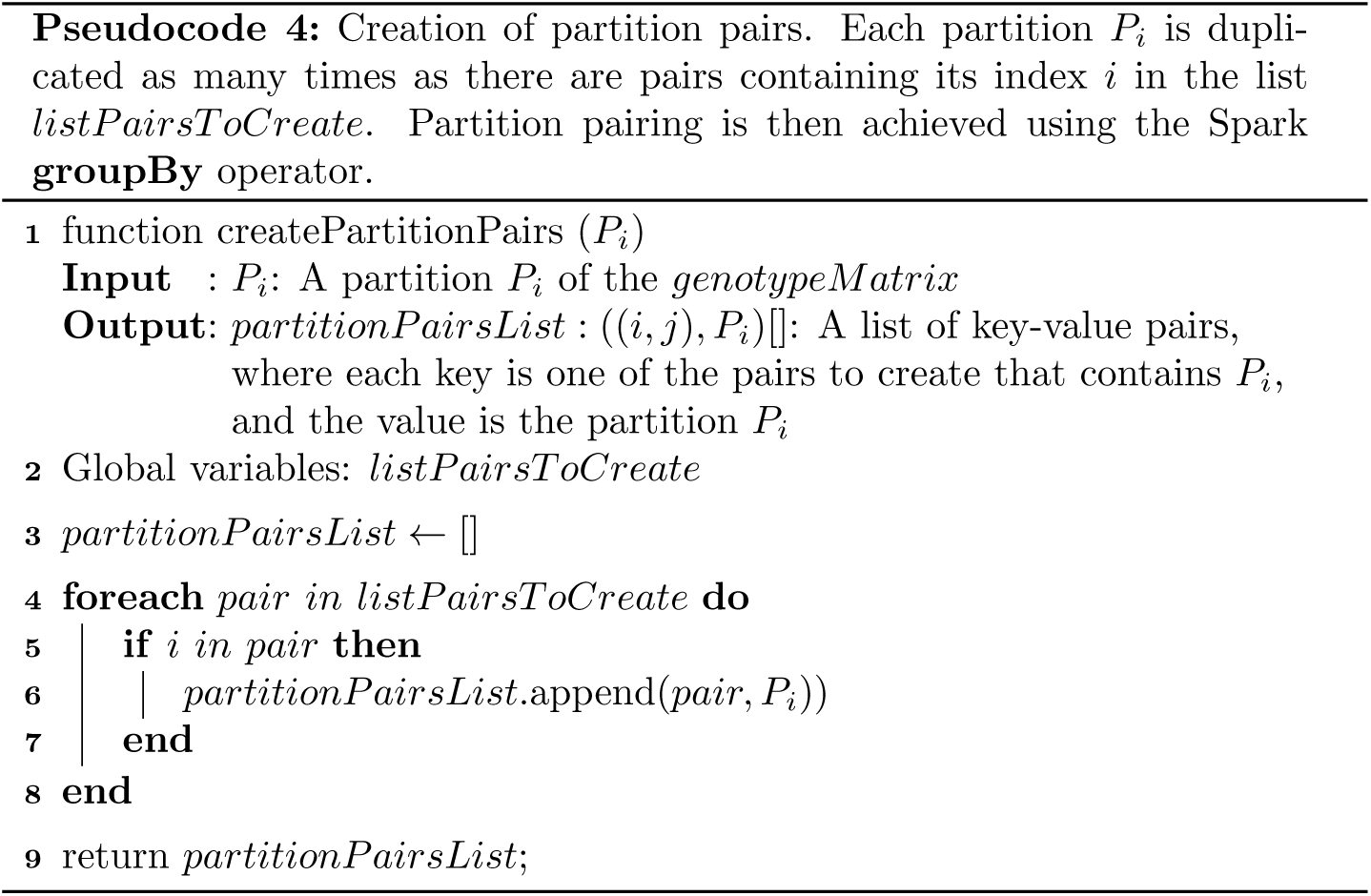

Index pairs are then grouped together, resulting in an RDD made of all the pairs of partitions ((*i*,*j*), (*P_i_*,*P_j_*)),*j* ≥ *i*. Each partition is finally scored, and scores are sorted and returned as a data frame. A summary of the sequence of transformation is given in Fig. 6.

**Figure 6:**
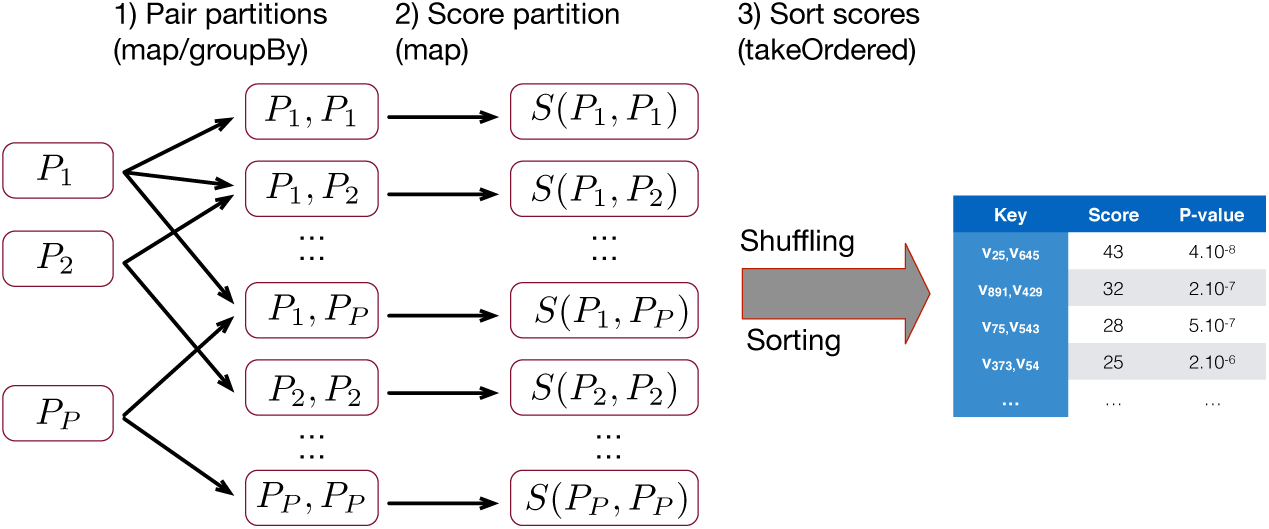
Pair of variant scoring: All *P*(*P*+1)/2 pairs of partitions are generated from the initial set of *P* partitions. Scoring is then performed for each partition pair, and scores are finally ranked and retrieved as a data frame.

The partition scoring now takes a pair of partition (*P_i_*,*P_j_*) as input, and returns the scores and p-values for possible pairs of variants in *P_i_* and *P_j_*.

##### 4) Extensions to larger genomic regions or *k*-way interactions

The proposed pipelines for gene and pair of variants scoring can be generalised to larger genomic regions (e.g., sets of genes) and *k*–way interactions (*k* > 2). The grouping of variants across multiple genes may be achieved by making the gene sets as keys, and by relying on the **groupBy** operator before the partition scoring, as was done for the gene scoring. The computation of scores involving *k*–way interactions (*k* > 2) is a generalisation of the *k* = 2 case. It can be achieved by modifying Pseudocode 4 so that *partitionPairsList* takes not only pairs of partitions, but any tuples of partition indices.

The generic scoring pipeline is summarised in Fig. 7, and consists in four main steps that sequentially performs variant grouping, partition interactions, scoring and sorting.

**Figure 7:**
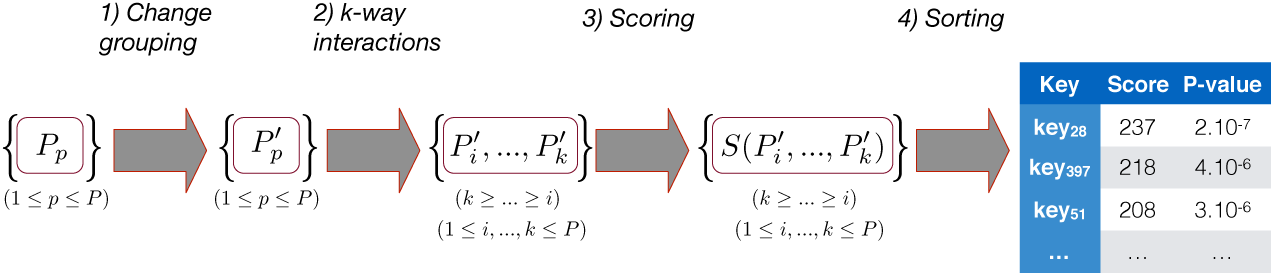
Scoring stage: Generic scoring pipeline. Steps 1 (change grouping) and 2 (k-way interactions) are optional preprocessing steps. The former allows to group variants across genes or sets of genes for aggregative scorings. The latter allows to assess interactions between genomic regions for multi-loci scorings. Steps 3 (scoring) and 4 (sorting) are common to all scoring workflows, and perform the actual scoring and ranking, respectively.

The *k*-way interaction step increases the number of partitions from *P* to an order *O* 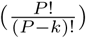, that is, all possible combinations of *k* partitions. For other steps, the number of partitions remains by default the same. It may however be changed programatically as an argument to the **groupBy** operator, or by explicitly using the **repartition** operator.

It is finally worth emphasising that all these steps can be expressed very concisely thanks to Spark’s grouping, mapping and sorting operators, which abstract the complexity of the distributed computing backend.

## User interface

DiGeST scoring pipeline may be called either from the command line, using a Spark submission script, or from a user-friendly Web front end (see online demo at http://bridgeiris.ulb.ac.be/digest). We detail both options below.

### Command line

The command line is the most direct way to run the DiGeST scoring pipeline. The parameters for an analysis are provided by means of a configuration file *jobArguments.conf*. The results are returned in two files: a CSV file containing the rankings, and a JSON file containing metadata about the analysis. This is illustrated in Fig. 8.

**Figure 8:**
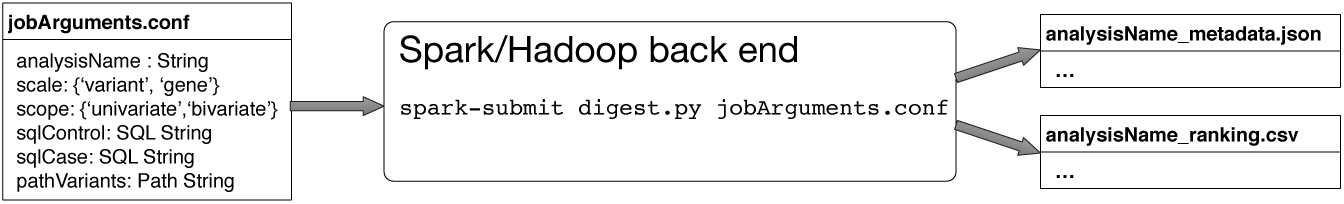
The scoring pipeline runs on a Spark/Hadoop back end. The scoring parameters are sent as arguments to the Spark job by means of a configuration file. The scoring results are returned in two files: A JSON file containing the metadata of the analysis, and a CSV file containing the variants/genes rankings together with their scores.

The configuration file must specify the following parameters:

- analysisName: The name for the analysis
- scale: Scale of the analysis, either ‘variant’ or ‘gene’
- scope: Scope of the analysis, either ‘univariate’ or ‘bivariate’
- sqlControl and sqlCase: The SQL queries for selecting variants for the control and case populations
- pathVariants: Path to the variant dataframe, in Parquet format.

The configuration file is passed as a parameter to the DiGeST Spark Python script ‘digest.py’, which returns two files after completion:

- analysisName_metadata.json : Contains the same as jobArguments.conf, plus the total runtime for the analysis
- analysisName_ranking.csv: A CSV file containing the variants/genes ranked according to their scores.

### Shiny R Web front end

The text files used as input and outputs of the DiGeST command line interface can be cumbersome to manage, especially for experiments involving a large number of genes, samples or filtering criteria. We therefore designed a Web application that allows on the one hand to create filters and groups, and generate the jobsArguments.conf, and on the other hand to explore the ranking results in a friendly way. The Web application is developed with Shiny R [60], and provides four main tools for interactively managing DiGeST data.

#### 1) Filtering tool

The filtering tool allows to create sets of filtering criteria for defining case and control populations. Filters are conditions of the form

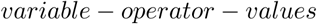

for example, *filters equal PASS*. Conditions can be combined and nested, allowing to express a large range of conditions on the variants to include. A snapshot of a filter that selects variants from the European samples of the 1000 Genomes Project, where the filtering quality is *PASS*, and SnpEff impact [13] is either high or moderate is given in Fig. 9.

**Figure 9:**
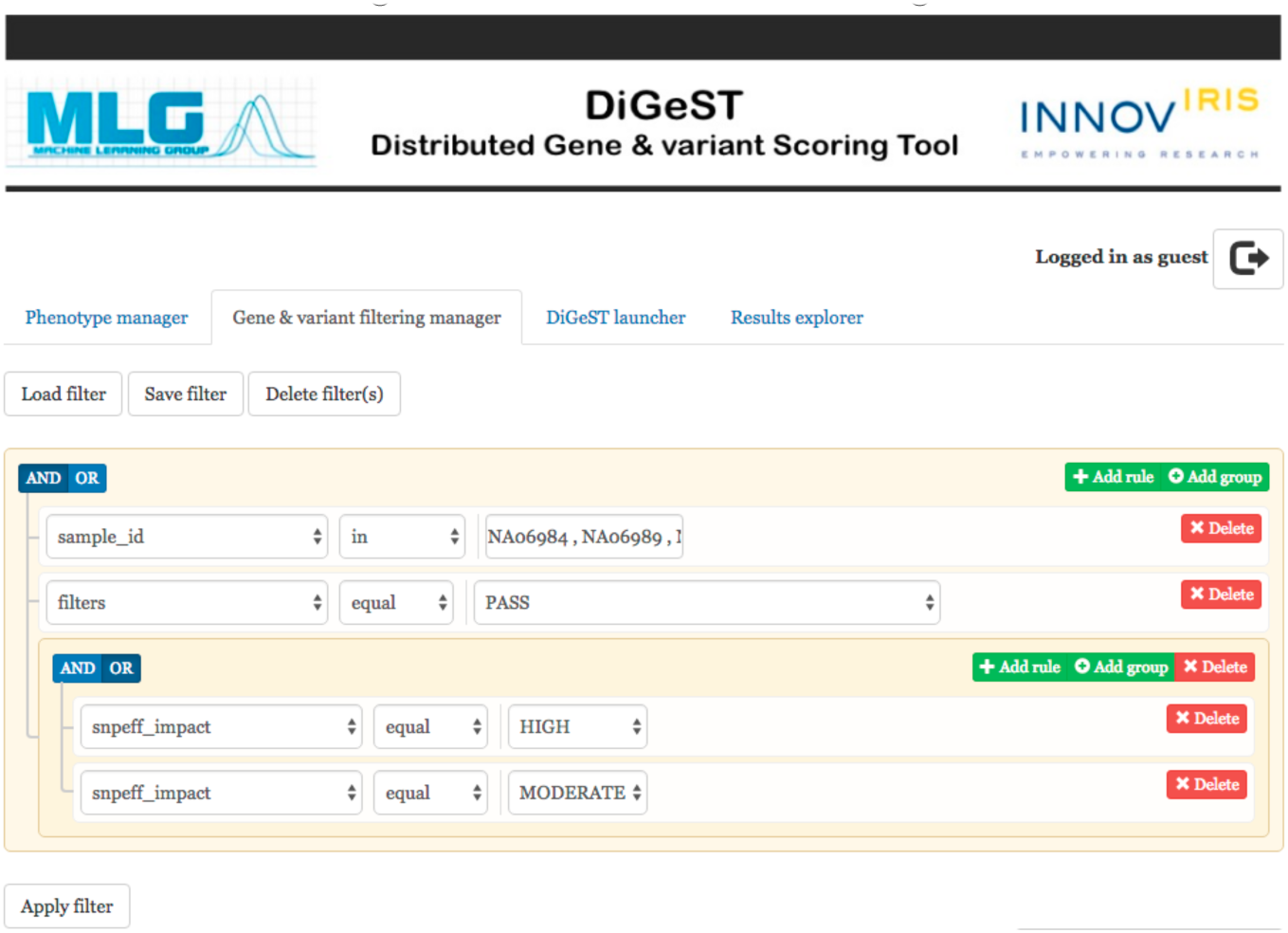
Web front-end: Filtering tool.

The set of variables proposed by the filtering tool are those included in the variant dataframe. In our design, we preprocessed the exome data from the 1000 Genomes Project using the Highlander filtering tool [24], and included 35 annotations fields from the 1000 Genomes Project [8], dbSNP [63], dbNSFP [38], and SnpEFF [13].

The filtering tool allows to save a given set of filtering conditions. The conditions are converted in a SQL syntax, that can either be used to feed the *jobsArgument.conf* file, or to interactively query the variant dataframe from the Parquet files (using the *Apply filter* button). In the latter case, a subset of 1000 variants are retrieved from the variant dataframe, and provided to the user as a table than can be either explored from the Web interface, or downloaded as a CSV. The query from Fig. 9, for example matches around 5.5 million variants. A snapshot of the table provided to the user is given in Fig. 10.

**Figure 10:**
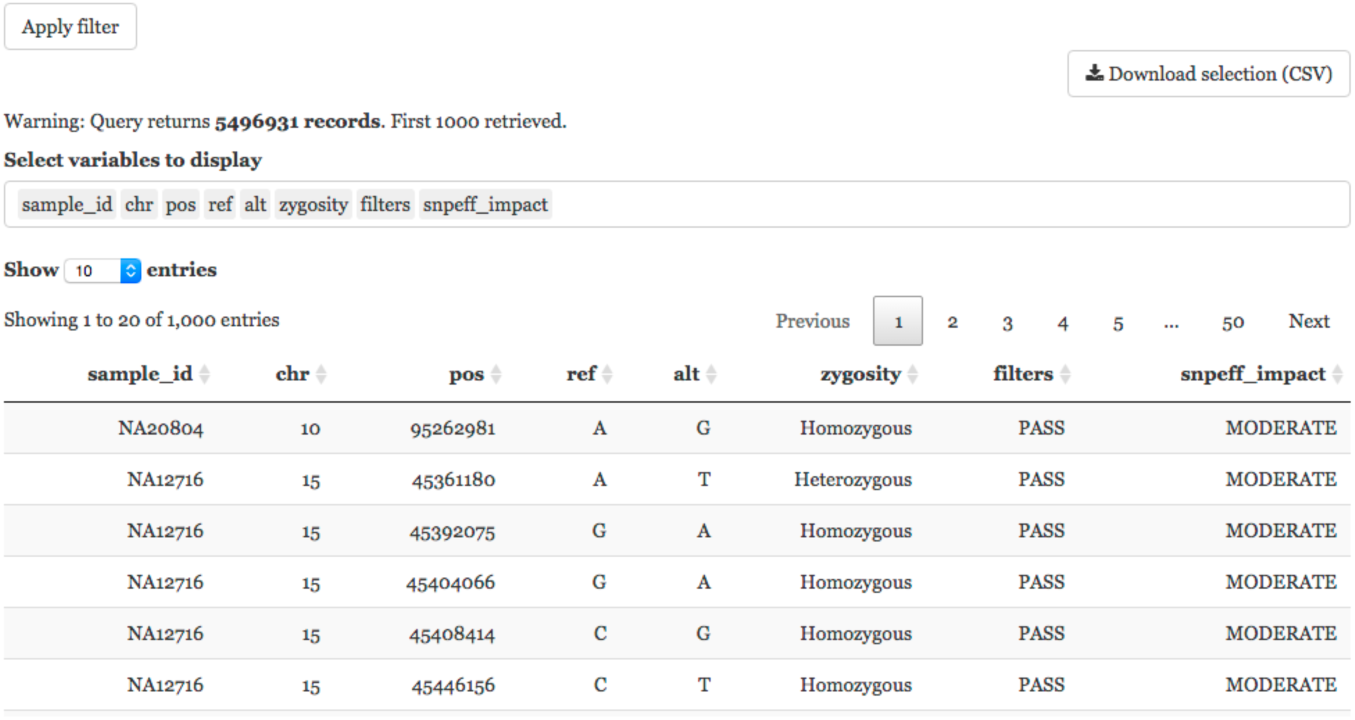
Web front-end: Variant table.

The filtering tool is available both for phenotypic and variant data, under the ‘Phenotype manager’ and ‘Gene and variant filtering manager’ (see Fig. 9), respectively.

#### 2) DiGeST launcher

The DiGeST launcher allows to create the *jobsArguments.conf* file, and start an analysis. Control and case groups are selected from the set of previously saved filters. The user may then select the scale (variant or gene) and scope (univariate or bivariate) for the scoring, and give a name to the analysis. The *jobsArguments.conf* file is created once pressing the start button, and provided to the Spark submission script.

#### 3) Results browser

The DiGeST pipeline returns the rankings in the form of a CSV file, whose rows contain the variant/gene or pair of variants/genes IDs, together with the scores and a set of metadata (p-values, number of variants, scores for individual groups). The results browser allows to open the CSV of an analysis, and to display the results in an interactive table where results may be reordered according to the columns. Depending on the scope of the analysis (variant, gene, pair of genes), the table provides the variant ID (in the form CHR:POS:REF:ALT), gene symbol (HGNC gene nomenclature [19]), or pair of variants IDs or genes symbols. A hyperlink connects gene symbols to their page on the OMIM (Online Mendelian Inheritance in Man) Web site [21]. An example of the result table is given in Fig. 12 for a digenic ranking.

**Figure 11:**
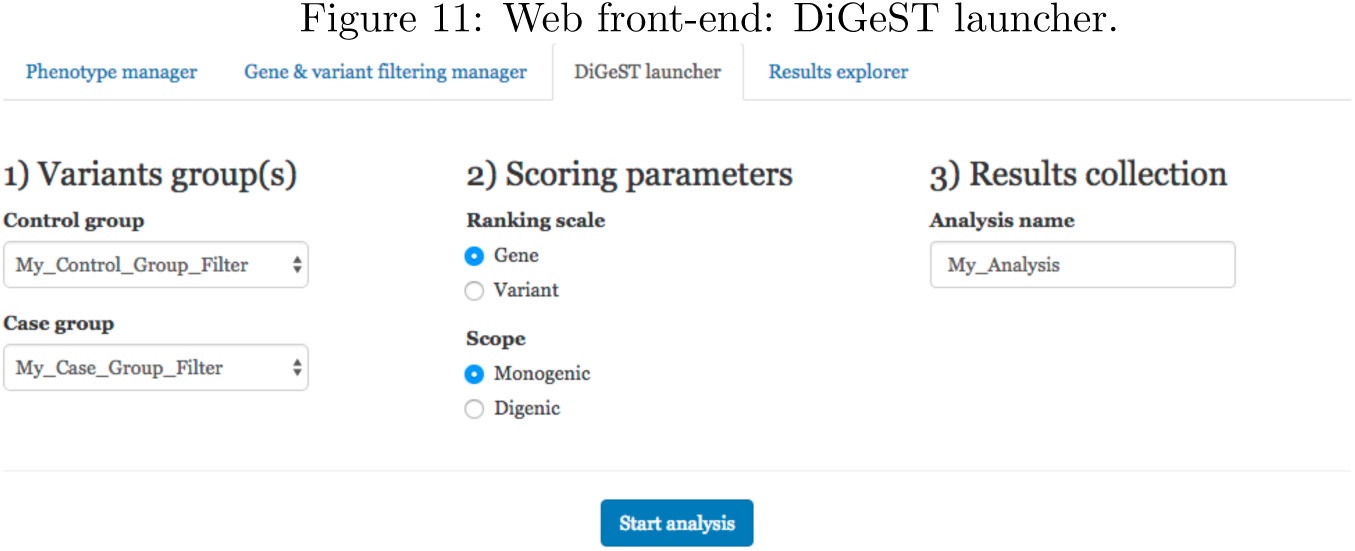
Web front-end: DiGeST launcher.

**Figure 12:**
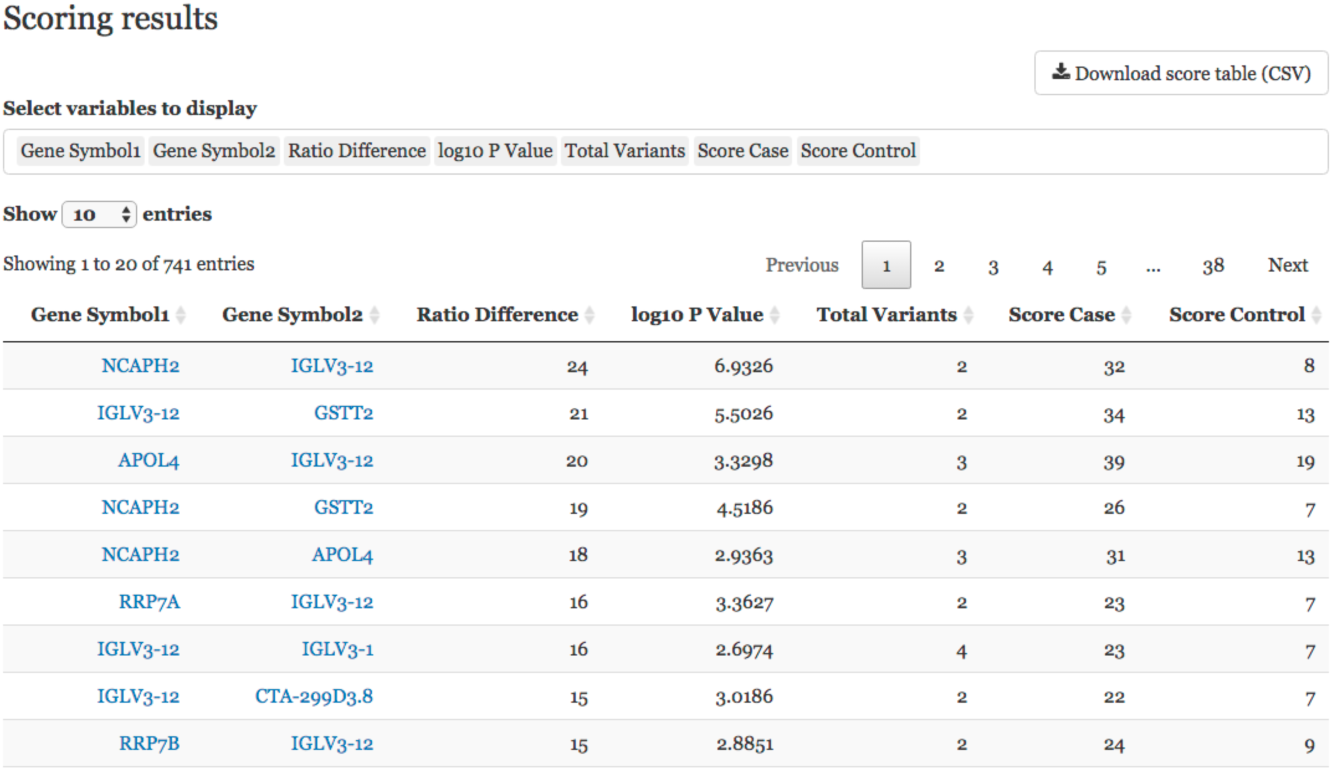
Web front-end: Results browser.

Clicking on an entry in the table provides a second table giving all the variants involved in a given scoring. For variant ranking, this retrieves details of a variant for all samples (control and case) involved in the analysis. For gene rankings, all variants belonging to the gene (or pair of genes) from all control and case samples are retrieved. That second table allows to analyse in details which samples, or variants, are effectively involved in the resulting scores.

#### 4) Pivot table

The Web user interface (UI) finally provides a pivot table tool, allowing to rearrange and visualise variant data in a variety of ways. The pivot table is available as a complementary widget to the filtering tool and results browser.

In the result browser, the pivot table is particularly useful to visualise how variants are distributed among samples of the case and control groups. An example is given in Fig. 13, where the number of variants in each gene of the first pair (IGLV3-12 and NCAPH2) is given for each sample of the two groups (European and Asian subset of samples from the 1000 genome project).

**Figure 13:**
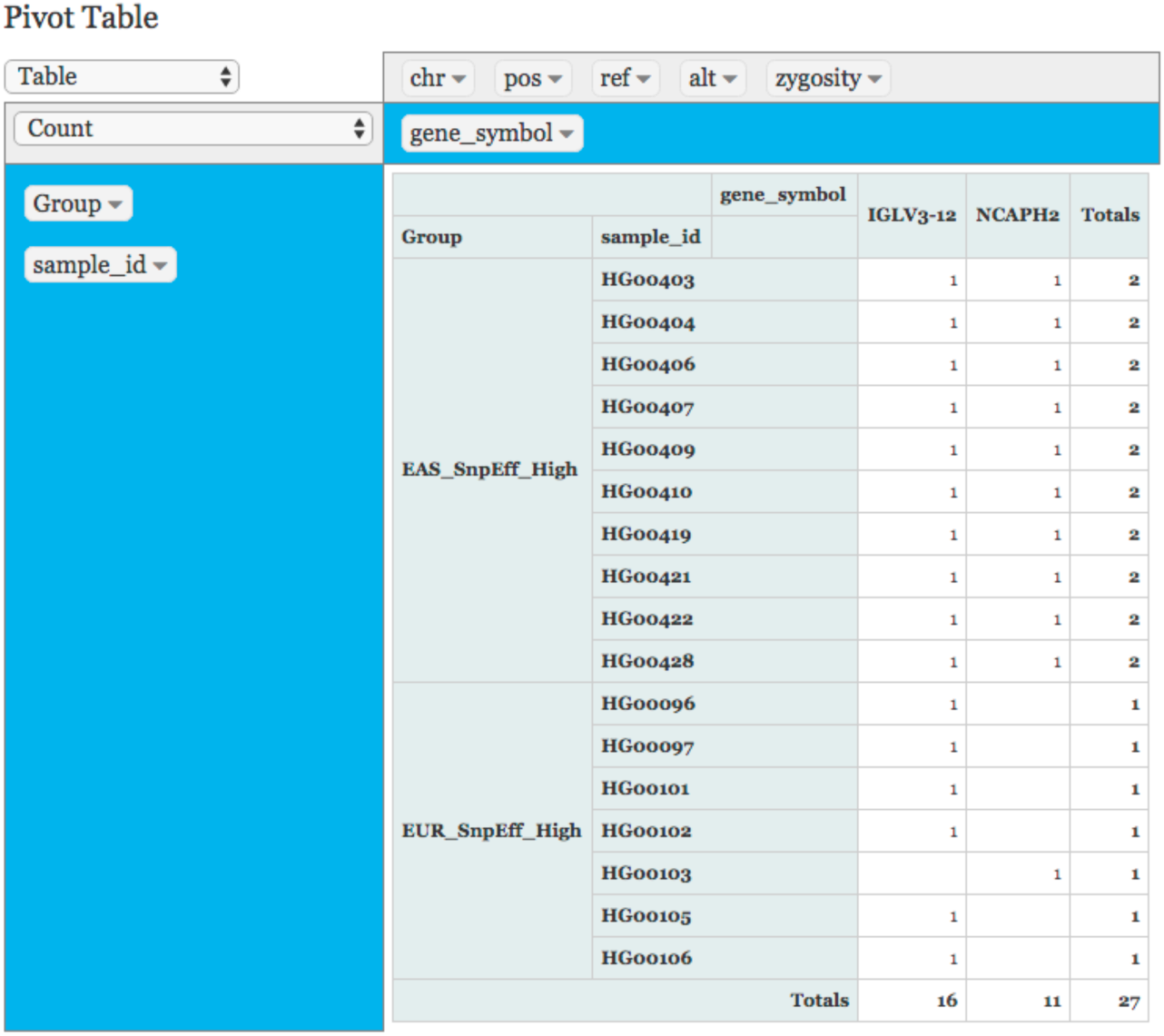
Web front-end: Pivot table.

## Results

We ran DiGeST on the 1000 Genomes Project phase 3 (1000GPp3) exome data [8], containing SNPs data for 2504 individuals, and a total of around 2.7 million variants. This section presents the data preparation and scalability results for univariate and bivariate rankings, both at the variant and gene scales.

### Data preparation

Exome variants were filtered and annotated using Highlander [24], resulting in a total of about 260 million annotated variants for all 2504 individuals (samples). Each sample has around 100000 variants, see Fig. 14, left. The second peak around 120000 variants per sample correspond to samples with African origin. The data covers 33168 genes (using UCSC hg19 genome assembly), and the median number of variants per gene is 47. A few genes (41) have more than 1000 variant per genes. The number of variants per genes is reported in Fig. 14, right (genes with more than 1000 variants are not represented for clarity reason). Data was converted to the Parquet format, and the database size on disk after conversion was slightly less than 40GB.

**Figure 14:**
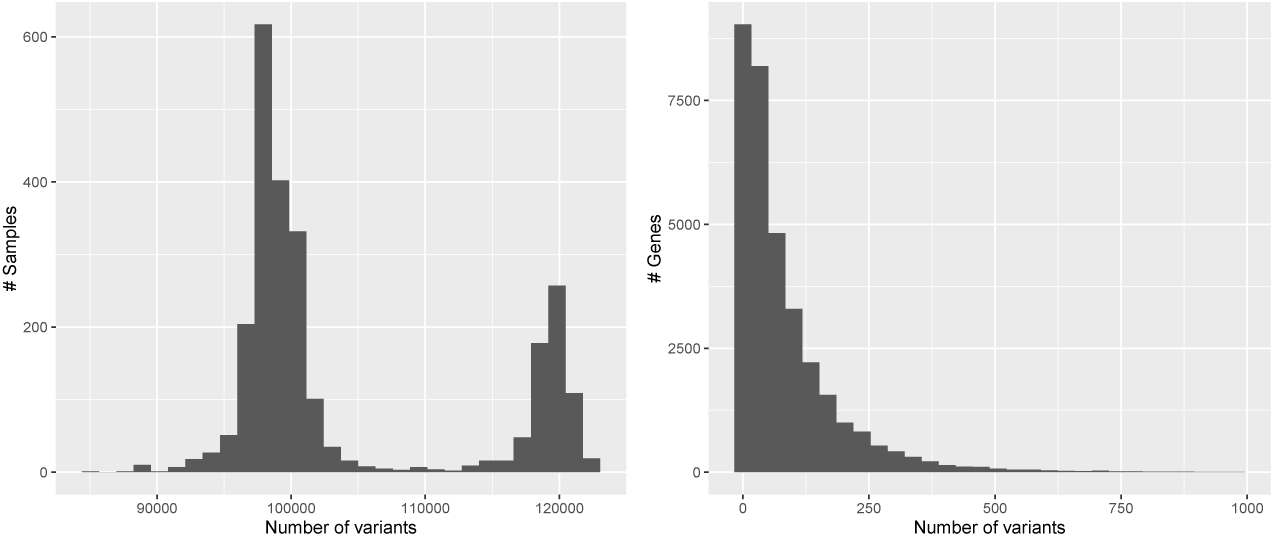
Histograms of the number of variants per sample (left) and per gene (right).

### Cluster back-end

We ran experiments on an in-house cluster consisting of 10 nodes, each with 24 cores, 128GB RAM, 4TB HDFS disk space, and 10Gb/s Ethernet connection. The Hadoop Yarn resource negociator system from Cloudera Hadoop Distribution 5.7.1 was used as the back-end for Spark 2.0. In all the experiments, Spark executors were allocated 2GB of RAM. The requested computing resources ranged from one executor, 2GB of RAM (single executor experiments) to one hundred executors and 200GB of RAM. All analyses were repeated five times, which was deemed sufficient given that computation times were very stable across runs.

### Variant rankings

We assessed the computation times for variant rankings by varying the number of variants from 10k to 2.8 millions, for a population of 2504 samples (1252 samples in the control and case populations). Scalability results are reported in Fig. 15, for a number of executors varying from 1 to 50. The left panel reports the genotype matrix creation times, and the right panel the corresponding variant scoring times.

**Figure 15:**
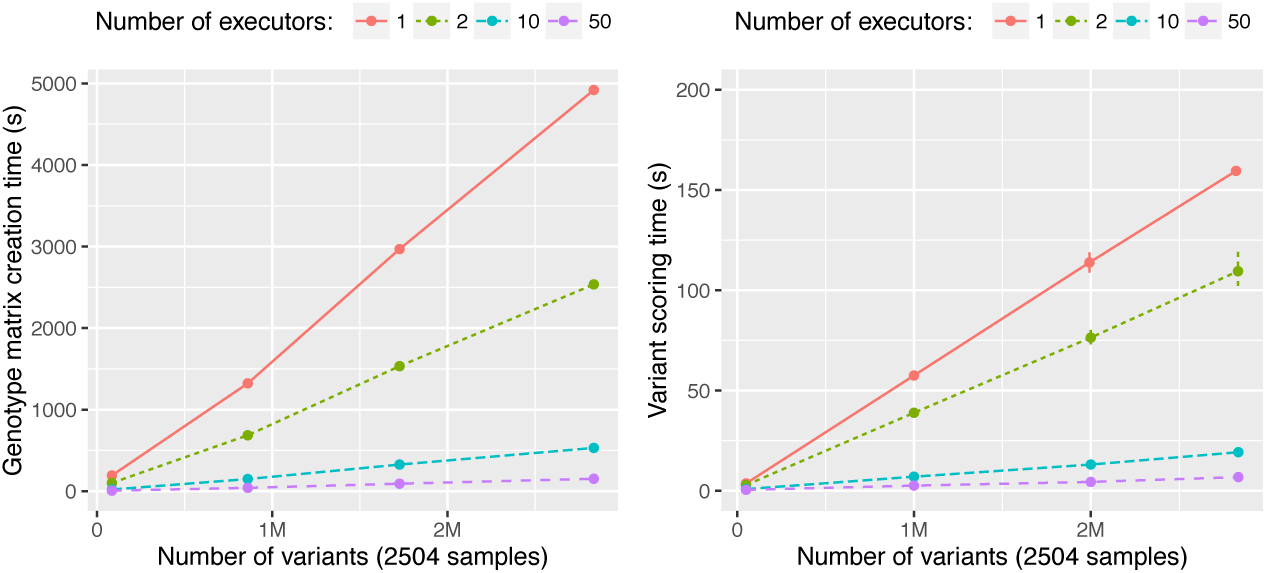
Scalability results for variant rankings. Execution times are reported separately for the genotype matrix creation times (left panel) and the variant scoring times (right panel).

The genotype matrix creation times were proportional to the number of variants included in the analysis (Fig. 15, left panel). The bottleneck for the creation time is the reading from the HDFS file system, which was on average of 8MB/s for our cluster. Given that the complete set of data was around 40GB, it took around 5000 seconds to create the whole genotype matrix with one executor. Thanks to the distributed file system, the loading times could be decreased by increasing the number of Spark executors. Loading times with 2 and 10 executors were reduced to 2500 and 530 seconds, providing almost proportional speed-ups of 1.9 and 9.2, respectively. Achievable speed-ups were however also bounded by the network bandwidth, which at a certain point becomes the main bottleneck. With 50 executors, the shuffling of data between executors reached its maximum capacity of 10Gb/s. The loading time was reduced was reduced to 150s, that is, a speed-up of only 30.

The computation times for scoring (Fig. 15, right panel) were also proportional to the number of variants. With one executor, the completion time for scoring all 2.7 million variants across the 2504 samples was of around 160s. The bottleneck were the disk access times, to load the 1.5GB sparse genotype matrix at a speed of again around 8MB/s. Increasing the number of executors to 10 and 50 reduced the computation times to 19s and 7s, providing speed-ups of 8.3 and 23, respectively. Given the relatively short computation times, Spark’s overhead becomes non negligible, therefore making these speedups sublinear in the number of executors. However, it is still remarkable that such speed-ups can be obtained given the short execution times of the task, and illustrate Spark’s ability to still provide speed-ups even for very short tasks.

Finally, it is worth noting that the standard deviations across all runs are very tight, on average less than one second. Only for one experiment (variant scoring, 2.7M variants, 2 executors) is the standard deviation noticeable on the graph (Fig. 15, right panel).

### Scalability of Gene rankings

The main difference between the gene and variant ranking pipelines is an added step that groups variants by genes after the genotype matrix is created. The genotype matrix creation times are therefore the same as in Fig. 15, left panel. We report below the processing times for the gene grouping step (Fig. 16, left panel), and the gene scoring step (Fig. 16, right panel) as the number of included genes increases from 1000 to 33000. As for the experiments on variant scorings, all 2504 samples were included (1252 in the control and case populations), and we assessed scalability by varying the number of executors from one to 50.

**Figure 16:**
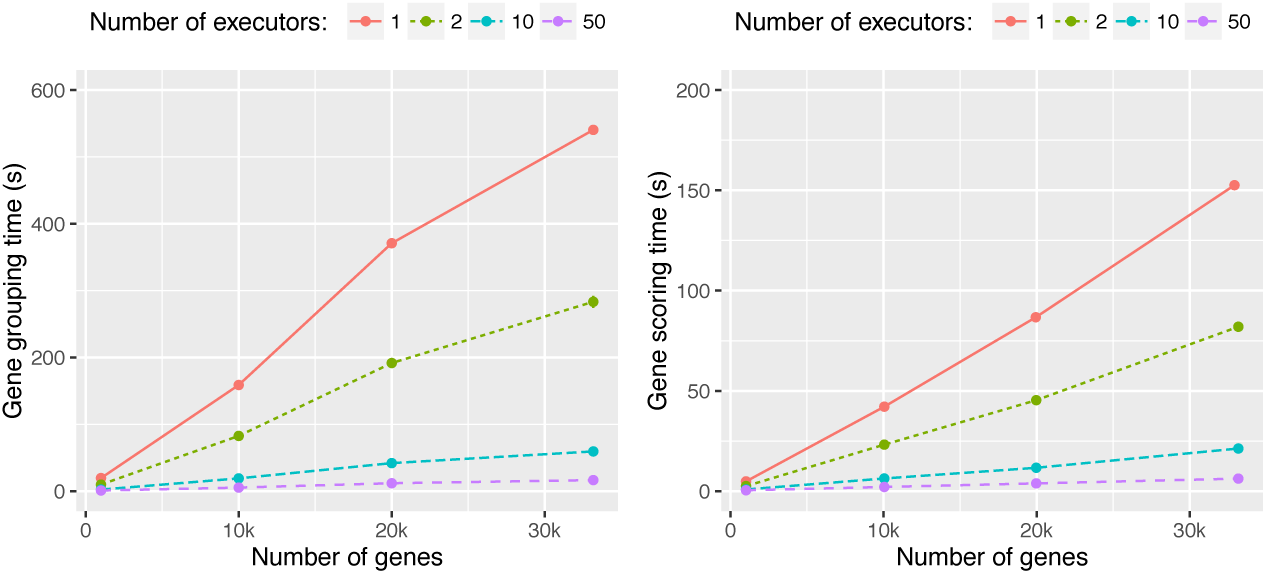
Scalability results for gene rankings. Execution times are reported separately for the genotype grouping times (left panel) and the gene scoring times (right panel).

The interpretation of the results is essentially similar to the scoring of variants: processing times follow a linear trend as the number of genes increases, and are bounded by disk access times. Hence, the scoring times are very comparable to those obtained for variant scoring. Grouping times took slightly longer (about twice the time), since shuffling occurs to group variants by genes. Processing times were also very stable across runs, with standard deviations less than a second, hence not visible on the graphs.

### Scalability of variant pair rankings

Scalability results are reported in Fig. 17, for a number of executors varying from 1 to 100. The left panel reports the computation times as the number of variants increases from 10k to 100k for a population size of 2504 samples, and the right panel reports the computation times as the population size increases from 100 to 2504 samples, for 25k variants. A timeout of 3600 seconds was set so that any run taking more than one hour was prematurely stopped.

**Figure 17:**
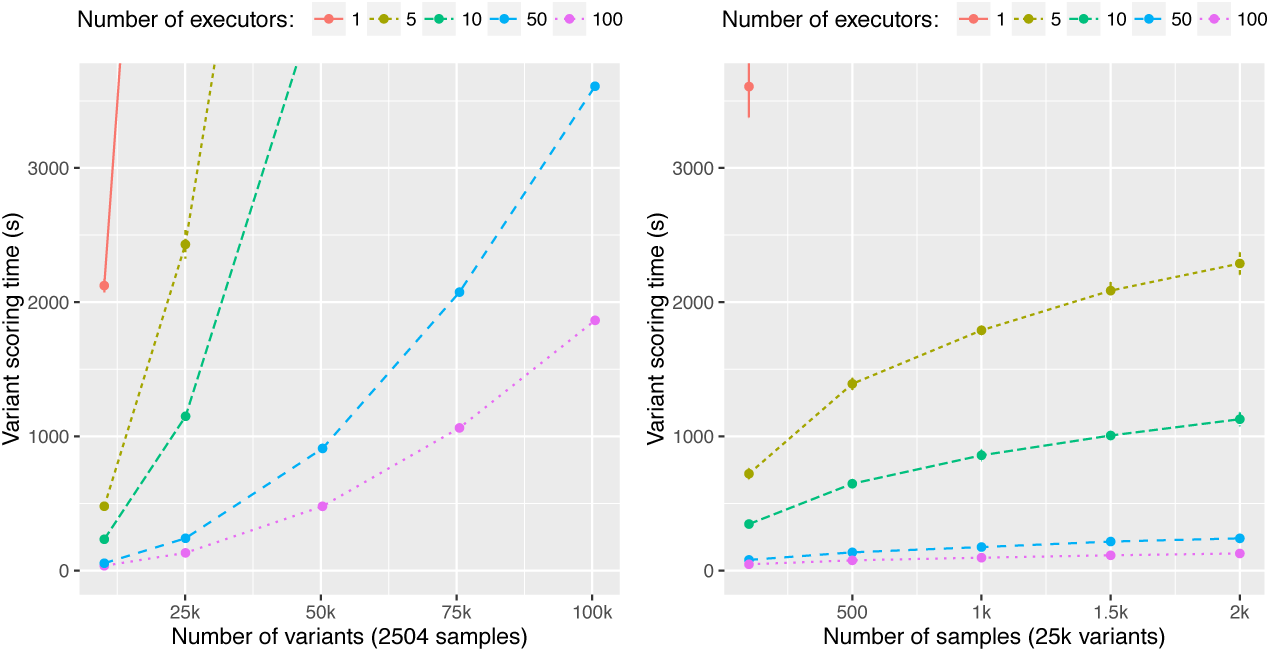
Scalability results for pairs of variants rankings. Execution times are reported separately for varying the number of variants (left panel), and varying the number of samples (right panel).

Computation times in the left panel follow a quadratic trend, that reflects the number of combinations 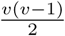 that are scored as the number of variants *v* is increased. For 10k variants (≈ 5 million pairs), the computation time for one executor is around 2120 seconds (with a standard deviation of 50 seconds), and decreases to 479 and 233 seconds with 5 and 10 executors (speed-ups of 4.4 and 9.1, respectively). Further increasing of the number of executors to 50 and 100 decreases the computation times to 54 and 36 seconds (speed-ups of 40 and 59, respectively).

Spark’s ability to fully use high numbers of executors becomes apparent when billions of scores are computed. Thus, the computation times for 100k variants (≈ 5 billion pairs) takes just one hour (3610±30 seconds) with 50 executors, and half an hour (1864±22 seconds) with 100 executors. With only one executor, computations take slightly more than 2 days (not reported on the chart for clarity reason). Speed-ups gained by using 50 and 100 executors are therefore of 48 and 95, respectively, which reflects the low overhead caused by Spark’s framework.

Finally, the trends observed in the right panel (increase of the sample size for a fixed number of variants) are sublinear, reflecting the sparsity property of the genotype matrix.

## Discussion and research perspectives

With DiGeST, we showed that Hadoop and Spark provide well-suited and complementary computing frameworks for performing genomic association studies. A compelling advantage of the frameworks is that most of the complexity related to the distributed infrastructure is abstracted by the HDFS storage system and the MapReduce programming model. In particular, the latter provides all the high-level operators required to group, filter, transform or sort variant data. This enables to express in a programmatically very concise way a wide variety of scoring schema, including aggregative or multi-loci scorings.

Our experimental results further illustrated the ability of both frameworks to efficiently distribute the loading and processing of large amounts of data, and to scale almost proportionally with the available computing resources. The main bottlenecks in execution times were found to be related to hardware limits, in particular disk access and network bandwidth. We plan to address in the future work a theoretical analysis of the tradeoffs between communication and parallelism, using models such as proposed in [61, 1].

Hadoop/Spark is designed to operate on commodity machines, and it should therefore be kept in mind that alternative computing frameworks such as High Performance Computing solutions, or those based on dedicated hardware such as GPUs [72, 71, 25, 18] and FPGAs [68, 20], still retain the edge in terms of raw computational capacity. Rather, the main benefits of Hadoop/Spark lie in its robustness to node failure, and its ability to provide an abstraction of the distributed infrastructure. Both these characteristics considerably speed up the development time of a distributed application.

Our work is, to the best of our knowledge, the first to provide an integrated Hadoop/Spark prototype for genomic association studies. The benefits of the Hadoop/Spark frameworks have however been recognised and successfully applied to a number of other bioinformatics tasks, such as genomic sequence mapping (CloudAligner [51], CloudBurst [62], SEAL [56]), genomic and RNA sequencing analysis (Crossbow [29], Myrna [28], Eoulsan [26]), sequence file management (Hadoop-BAM [52], SeqWare [11], GATK [44]), or phylogenetic analysis (MrsRF [43], Nephele [7]), see [33, 73, 54] for comprehensive reviews. In this ecosystem of tools and applications, an important research direction will consist in providing better integration and interoperability.

More specifically focusing on genomic association studies, we aim at extending DiGeST with a wider range of scoring functions. DiGeST currently only implements the basic association test for a disease trait by comparing allele frequencies between cases and controls, and applying a Fisher’s test. Extension to other standard GAS tests such as the Cochran-Armitage test and Hotelling’s T(2) statistic [58] are expected to require only slight variations in the partition scoring step of DiGeST pipeline. Extensions to aggregative tests such as burden tests (e.g., CMC [35], or CAST [48], WSS [40], KBAC [37]) or variance-component tests (e.g., SKAT [70]) could on the other hand be implemented by relying on the variant grouping feature of DiGeST.

Beyond scoring methods based on statistical tests, advanced computational techniques based on Machine Learning (ML) have more recently been showed to provide significant improvements to variant or gene prioritisation tasks [27, 64, 59, 47, 69, 14, 3]. A more general avenue for our research aims at investigating the integration of such techniques within the DiGeST pipeline.

## Conclusions

In the landscape of currently available open source distributed computing frameworks, the work presented in this article supports that Hadoop and Spark bring effective complementary solutions for DNA sequencing association studies. On the one hand, Hadoop is a robust and now well-established framework for fault-tolerant distributed data storage and resource management system. On the other hand, Spark provides a fast and light-weight computing engine, with a rich range of high-level functions for filtering, grouping, and transforming data in an efficient way.

Relying on Hadoop/Spark, we developed DiGeST, a distributed gene and variant scoring tool, and showed its ability to provide scalability to DNA sequencing case/control studies. We coupled this distributed back-end with a user-friendly Web front-end based on R Shiny, allowing users to easily filter variant data, and explore scoring results. All tools are made open-source, and can be reused efficiently thanks to virtualisation scripts made available from the project web site.

## References

[1] F. Afrati and J. Ullman. Matching bounds for the all-pairs mapreduce problem. In Proceedings of the 17th International Database Engineering & Applications Symposium, pages 3–4. ACM, 2013.

[2] G. Bhatia, V. Bansal, O. Harismendy, N. J. Schork, E. J. Topol, K. Frazer, and V. Bafna. A covering method for detecting genetic associations between rare variants and common phenotypes. PLoS Comput Biol, 6(10):e1000954, 2010.

[3] V. Botta, G. Louppe, P. Geurts, and L. Wehenkel. Exploiting snp correlations within random forest for genome-wide association studies. PloS one, 9(4):e93379, 2014.

[4] R. M. Cantor, K. Lange, and J. S. Sinsheimer. Prioritizing gwas results: a review of statistical methods and recommendations for their application. The American Journal of Human Genetics, 86(1): 6–22, 2010.

[5] P. Cingolani, V. Patel, M. Coon, T. Nguyen, S. Land, D. Ruden, and X. Lu. Using drosophila melanogaster as a model for genotoxic chemical mutational studies with a new program, snpsift. Frontiers in Genetics, 3, 2012.

[6] E. T. Cirulli and D. B. Goldstein. Uncovering the roles of rare variants in common disease through whole-genome sequencing. Nature Reviews Genetics, 11(6): 415–425, 2010.

[7] M. E. Colosimo, M. W. Peterson, S. Mardis, and L. Hirschman. Nephele: genotyping via complete composition vectors and mapreduce. Source code for biology and medicine, 6(1):1, 2011.

[8] G. P. Consortium et al. A global reference for human genetic variation. Nature, 526(7571): 68–74, 2015.

[9] H. J. Cordell. Detecting gene–gene interactions that underlie human diseases. Nature Reviews Genetics, 10(6): 392–404, 2009.

[10] H. J. Cordell and D. G. Clayton. Genetic association studies. The Lancet, 366(9491): 1121–1131, 2005.

[11] B. D O’Connor, B. Merriman, and S. F. Nelson. Seqware query engine: storing and searching sequence data in the cloud. BMC bioinformatics, 11(12):1, 2010.

[12] P. Danecek, A. Auton, G. Abecasis, C. A. Albers, E. Banks, M. A. DePristo, R. E. Handsaker, G. Lunter, G. T. Marth, S. T. Sherry, et al. The variant call format and vcftools. Bioinformatics, 27(15): 2156–2158, 2011.

[13] G. De Baets, J. Van Durme, J. Reumers, S. Maurer-Stroh, P. Vanhee, J. Dopazo, J. Schymkowitz, and F. Rousseau. Snpeffect 4.0: on-line prediction of molecular and structural effects of protein-coding variants. Nucleic Acids Research, page gkr996, 2011.

[14] S. A. Gagliano, M. R. Barnes, M. E. Weale, and J. Knight. A bayesian method to incorporate hundreds of functional characteristics with association evidence to improve variant prioritization. PloS one, 9(5):e98122, 2014.

[15] A. M. Gazzo, D. Daneels, E. Cilia, M. Bonduelle, M. Abramowicz, S. Van Dooren, G. Smits, and T. Lenaerts. Dida: A curated and annotated digenic diseases database. Nucleic acids research, 44(D1):D900–D907, 2016.

[16] R. C. Gentleman, V. J. Carey, D. M. Bates, B. Bolstad, M. Dettling, S. Dudoit, B. Ellis, L. Gautier, Y. Ge, J. Gentry, et al. Bioconductor: open software development for computational biology and bioinformatics. Genome biology, 5(10):1, 2004.

[17] V. Geoffroy, C. Pizot, C. Redin, A. Piton, N. Vasli, C. Stoetzel, A. Blavier, J. Laporte, and J. Muller. Varank: a simple and powerful tool for ranking genetic variants. PeerJ, 3:e796, 2015.

[18] B. Goudey, D. Rawlinson, Q. Wang, F. Shi, H. Ferra, R. M. Campbell, L. Stern, M. T. Inouye, C. S. Ong, and A. Kowalczyk. Gwis-model-free, fast and exhaustive search for epistatic interactions in case-control gwas. BMC genomics, 14(3):1, 2013.

[19] K. A. Gray, B. Yates, R. L. Seal, M. W. Wright, and E. A. Bruford. Genenames. org: the hgnc resources in 2015. Nucleic acids research, page gku1071, 2014.

[20] S. Gundlach, J. C. Kässens, and L. Wienbrandt. Genome-wide association interaction studies with mb-mdr and maxt multiple testing correction on fpgas. Procedia Computer Science, 80:639–649, 2016.

[21] A. Hamosh, A. F. Scott, J. S. Amberger, C. A. Bocchini, and V. A. McKusick. Online mendelian inheritance in man (omim), a knowledgebase of human genes and genetic disorders. Nucleic acids research, 33(suppl 1):D514–D517, 2005.

[22] F. Han and W. Pan. A data-adaptive sum test for disease association with multiple common or rare variants. Human heredity, 70(1): 42–54, 2010.

[23] R. Helaers. Assotester: R package with statistical tests and methods for genetic association studies with emphasis on rare variants and binary (dichotomous) traits. In preparation.

[24] R. Helaers and M. Vikkula. Highlander: variant filtering made easy. Submitted.

[25] G. Hemani, A. Theocharidis, W. Wei, and C. Haley. Epigpu: exhaustive pairwise epistasis scans parallelized on consumer level graphics cards. Bioinformatics, 27(11): 1462–1465, 2011.

[26] L. Jourdren, M. Bernard, M.-A. Dillies, and S. Le Crom. Eoulsan: a cloud computing-based framework facilitating high throughput sequencing analyses. Bioinformatics, 28(11): 1542–1543, 2012.

[27] M. Kircher, D. M. Witten, P. Jain, B. J. O’Roak, G. M. Cooper, and J. Shendure. A general framework for estimating the relative pathogenicity of human genetic variants. Nature genetics, 46(3):310, 2014.

[28] B. Langmead, K. D. Hansen, and J. T. Leek. Cloud-scale rna-sequencing differential expression analysis with myrna. Genome biology, 11(8):1, 2010.

[29] B. Langmead, M. C. Schatz, J. Lin, M. Pop, and S. L. Salzberg. Searching for snps with cloud computing. Genome biology, 10(11):1, 2009.

[30] Y.-A. Le Borgne. Digest: Distributed gene and variant scoring tool.

[31] I.-H. Lee, K. Lee, M. Hsing, Y. Choe, J.-H. Park, S. H. Kim, J. M. Bohn, M. B. Neu, K.-B. Hwang, R. C. Green, et al. Prioritizing disease-linked variants, genes, and pathways with an interactive whole-genome analysis pipeline. Human mutation, 35(5): 537–547, 2014.

[32] S. Lee, G. R. Abecasis, M. Boehnke, and X. Lin. Rare-variant association analysis: study designs and statistical tests. The American Journal of Human Genetics, 95(1): 5–23, 2014.

[33] S. H. Lelieveld, J. A. Veltman, and C. Gilissen. Novel bioinformatic developments for exome sequencing. Human genetics, pages 1–12, 2016.

[34] C. M. Lewis and J. Knight. Introduction to genetic association studies. Cold Spring Harbor Protocols, 2012(3):pdb–top068163, 2012.

[35] B. Li and S. M. Leal. Methods for detecting associations with rare variants for common diseases: application to analysis of sequence data. The American Journal of Human Genetics, 83(3): 311–321, 2008.

[36] E. T. Lim, S. Raychaudhuri, S. J. Sanders, C. Stevens, A. Sabo, D. G. MacArthur, B. M. Neale, A. Kirby, D. M. Ruderfer, M. Fromer, et al. Rare complete knockouts in humans: population distribution and significant role in autism spectrum disorders. Neuron, 77(2): 235–242, 2013.

[37] D. J. Liu and S. M. Leal. A novel adaptive method for the analysis of next-generation sequencing data to detect complex trait associations with rare variants due to gene main effects and interactions. PLoS Genet, 6(10):e1001156, 2010.

[38] X. Liu, X. Jian, and E. Boerwinkle. dbnsfp v2. 0: a database of human nonsynonymous snvs and their functional predictions and annotations. Human mutation, 34(9):E2393–E2402, 2013.

[39] T. F. Mackay. Epistasis and quantitative traits: using model organisms to study gene-gene interactions. Nature Reviews Genetics, 15(1): 22–33, 2014.

[40] B. E. Madsen and S. R. Browning. A groupwise association test for rare mutations using a weighted sum statistic. PLoS Genet, 5(2):e1000384, 2009.

[41] J. Marchini, P. Donnelly, and L. R. Cardon. Genome-wide strategies for detecting multiple loci that influence complex diseases. Nature genetics, 37(4): 413–417, 2005.

[42] M. Massie, F. Nothaft, C. Hartl, C. Kozanitis, A. Schumacher, A. D. Joseph, and D. A. Patterson. Adam: Genomics formats and processing patterns for cloud scale computing. University of California, Berkeley Technical Report, No. UCB/EECS-2013, 207, 2013.

[43] S. J. Matthews and T. L. Williams. Mrsrf: an efficient mapreduce algorithm for analyzing large collections of evolutionary trees. BMC bioinformatics, 11(1):11, 2010.

[44] A. McKenna, M. Hanna, E. Banks, A. Sivachenko, K. Cibulskis, A. Kernytsky, K. Garimella, D. Altshuler, S. Gabriel, M. Daly, et al. The genome analysis toolkit: a mapreduce framework for analyzing next-generation dna sequencing data. Genome research, 20(9): 1297–1303, 2010.

[45] B. A. McKinney, D. M. Reif, M. D. Ritchie, and J. H. Moore. Machine learning for detecting gene-gene interactions. Applied bioinformatics, 5(2): 77–88, 2006.

[46] S. Melnik, A. Gubarev, J. J. Long, G. Romer, S. Shivakumar, M. Tolton, and T. Vassilakis. Dremel: Interactive analysis of web-scale datasets. In Proc. of the 36th Int’l Conf on Very Large Data Bases, pages 330–339, 2010.

[47] F. Mordelet and J.-P. Vert. Prodige: Prioritization of disease genes with multitask machine learning from positive and unlabeled examples. BMC bioinformatics, 12(1):1, 2011.

[48] S. Morgenthaler and W. G. Thilly. A strategy to discover genes that carry multi-allelic or mono-allelic risk for common diseases: a cohort allelic sums test (cast). Mutation Research/Fundamental and Molecular Mechanisms of Mutagenesis, 615(1): 28–56, 2007.

[49] A. P. Morris and E. Zeggini. An evaluation of statistical approaches to rare variant analysis in genetic association studies. Genetic epidemiology, 34(2): 188–193, 2010.

[50] B. M. Neale, M. A. Rivas, B. F. Voight, D. Altshuler, B. Devlin, M. Orho-Melander, S. Kathiresan, S. M. Purcell, K. Roeder, and M. J. Daly. Testing for an unusual distribution of rare variants. PLoS Genet, 7(3):e1001322, 2011.

[51] T. Nguyen, W. Shi, and D. Ruden. Cloudaligner: A fast and full-featured mapreduce based tool for sequence mapping. BMC research notes, 4(1):1, 2011.

[52] M. Niemenmaa, A. Kallio, A. Schumacher, P. Klemelä, E. Korpelainen, and K. Heljanko. Hadoop-bam: directly manipulating next generation sequencing data in the cloud. Bioinformatics, 28(6): 876–877, 2012.

[53] S. Oh, J. Lee, M.-S. Kwon, B. Weir, K. Ha, and T. Park. A novel method to identify high order gene-gene interactions in genome-wide association studies: Gene-based mdr. BMC bioinformatics, 13(9):1, 2012.

[54] A. O’Driscoll, J. Daugelaite, and R. D. Sleator. ‘big data’, hadoop and cloud computing in genomics. Journal of biomedical informatics, 46(5): 774–781, 2013.

[55] S. Pabinger, A. Dander, M. Fischer, R. Snajder, M. Sperk, M. Efremova, B. Krabichler, M. R. Speicher, J. Zschocke, and Z. Trajanoski. A survey of tools for variant analysis of next-generation genome sequencing data. Briefings in bioinformatics, 15(2): 256–278, 2014.

[56] L. Pireddu, S. Leo, and G. Zanetti. Seal: a distributed short read mapping and duplicate removal tool. Bioinformatics, 27(15): 2159–2160, 2011.

[57] A. L. Price, G. V. Kryukov, P. I. de Bakker, S. M. Purcell, J. Staples, L.-J. Wei, and S. R. Sunyaev. Pooled association tests for rare variants in exon-resequencing studies. The American Journal of Human Genetics, 86(6): 832–838, 2010.

[58] S. Purcell, B. Neale, K. Todd-Brown, L. Thomas, M. A. Ferreira, D. Bender, J. Maller, P. Sklar, P. I. De Bakker, M. J. Daly, et al. Plink: a tool set for whole-genome association and population-based linkage analyses. The American Journal of Human Genetics, 81(3): 559–575, 2007.

[59] G. R. Ritchie, I. Dunham, E. Zeggini, and P. Flicek. Functional annotation of noncoding sequence variants. Nature methods, 11(3): 294–296, 2014.

[60] RStudio, Inc. Easy web applications in R., 2013. URL: http://www.rstudio.com/shiny/.

[61] A. D. Sarma, F. N. Afrati, S. Salihoglu, and J. D. Ullman. Upper and lower bounds on the cost of a map-reduce computation. In Proceedings of the VLDB Endowment, volume 6, pages 277–288. VLDB Endowment, 2013.

[62] M. C. Schatz. Cloudburst: highly sensitive read mapping with mapreduce. Bioinformatics, 25(11): 1363–1369, 2009.

[63] S. T. Sherry, M.-H. Ward, M. Kholodov, J. Baker, L. Phan, E. M. Smigielski, and K. Sirotkin. dbsnp: the ncbi database of genetic variation. Nucleic acids research, 29(1): 308–311, 2001.

[64] A. Sifrim, D. Popovic, L.-C. Tranchevent, A. Ardeshirdavani, R. Sakai, P. Konings, J. R. Vermeesch, J. Aerts, B. De Moor, and Y. Moreau. extasy: variant prioritization by genomic data fusion. Nature methods, 10(11): 1083–1084, 2013.

[65] N. O. Stitziel, A. Kiezun, and S. Sunyaev. Computational and statistical approaches to analyzing variants identified by exome sequencing. Genome biology, 12(9):1, 2011.

[66] X. Wan, C. Yang, Q. Yang, H. Xue, X. Fan, N. L. Tang, and W. Yu. Boost: A fast approach to detecting gene-gene interactions in genome-wide case-control studies. The American Journal of Human Genetics, 87(3): 325–340, 2010.

[67] G. T. Wang, B. Peng, and S. M. Leal. Variant association tools for quality control and analysis of large-scale sequence and genotyping array data. The American Journal of Human Genetics, 94(5): 770–783, 2014.

[68] L. Wienbrandt, J. C. Kässens, J. González-Domínguez, B. Schmidt, D. Ellinghaus, and M. Schimmler. Fpga-based acceleration of detecting statistical epistasis in gwas. Procedia Computer Science, 29:220–230, 2014.

[69] M. Wu, J. Wu, T. Chen, and R. Jiang. Prioritization of nonsynonymous single nucleotide variants for exome sequencing studies via integrative learning on multiple genomic data. Scientific reports, 5, 2015.

[70] M. C. Wu, S. Lee, T. Cai, Y. Li, M. Boehnke, and X. Lin. Rare-variant association testing for sequencing data with the sequence kernel association test. The American Journal of Human Genetics, 89(1): 82–93, 2011.

[71] G. Yang, W. Jiang, Q. Yang, and W. Yu. Pboost: A gpu based tool for parallel permutation tests in genome-wide association studies. Bioinformatics, page btu840, 2014.

[72] L. S. Yung, C. Yang, X. Wan, and W. Yu. Gboost: a gpu-based tool for detecting gene–gene interactions in genome–wide case control studies. Bioinformatics, 27(9): 1309–1310, 2011.

[73] Q. Zou, X.-B. Li, W.-R. Jiang, Z.-Y. Lin, G.-L. Li, and K. Chen. Survey of mapreduce frame operation in bioinformatics. Briefings in bioinformatics, page bbs088, 2013.

